# Transcriptomic Imputation of Bipolar Disorder and Bipolar subtypes reveals 29 novel associated genes

**DOI:** 10.1101/222786

**Authors:** Laura M. Huckins, Amanda Dobbyn, Whitney McFadden, Weiqing Wang, Douglas M. Ruderfer, Gabriel Hoffman, Veera Rajagopal, Hoang T. Nguyen, Panos Roussos, Menachem Fromer, Robin Kramer, Enrico Domenci, Eric Gamazon, CommonMind Consortium, the Bipolar Disorder Working Group of the Psychiatric Genomics Consortium, iPSYCH Consortium, Ditte Demontis, Anders Børglum, Bernie Devlin, Solveig K. Sieberts, Nancy Cox, Hae Kyung Im, Pamela Sklar, Eli A. Stahl

## Abstract

Bipolar disorder is a complex neuropsychiatric disorder presenting with episodic mood disturbances. In this study we use a transcriptomic imputation approach to identify novel genes and pathways associated with bipolar disorder, as well as three diagnostically and genetically distinct subtypes. Transcriptomic imputation approaches leverage well-curated and publicly available eQTL reference panels to create gene-expression prediction models, which may then be applied to “impute” genetically regulated gene expression (GREX) in large GWAS datasets. By testing for association between phenotype and GREX, rather than genotype, we hope to identify more biologically interpretable associations, and thus elucidate more of the genetic architecture of bipolar disorder.

We applied GREX prediction models for 13 brain regions (derived from CommonMind Consortium and GTEx eQTL reference panels) to 21,488 bipolar cases and 54,303 matched controls, constituting the largest transcriptomic imputation study of bipolar disorder (BPD) to date. Additionally, we analyzed three specific BPD subtypes, including 14,938 individuals with subtype 1 (BD-I), 3,543 individuals with subtype 2 (BD-II), and 1,500 individuals with schizoaffective subtype (SAB).

We identified 125 gene-tissue associations with BPD, of which 53 represent independent associations after FINEMAP analysis. 29/53 associations were novel; i.e., did not lie within 1Mb of a locus identified in the recent PGC-BD GWAS. We identified 37 independent BD-I gene-tissue associations (10 novel), 2 BD-II associations, and 2 SAB associations. Our BPD, BD-I and BD-II associations were significantly more likely to be differentially expressed in post-mortem brain tissue of BPD, BD-I and BD-II cases than we might expect by chance. Together with our pathway analysis, our results support long-standing hypotheses about bipolar disorder risk, including a role for oxidative stress and mitochondrial dysfunction, the post-synaptic density, and an enrichment of circadian rhythm and clock genes within our results.

## Introduction

Bipolar disorder (BPD) is a serious episodic neuropsychiatric disorder presenting with extreme elation, or mania, and severe depressive states^1^. In tandem, individuals with bipolar often experience disturbances in thinking and behavior, as well as psychotic features such as delusions and hallucinations^1^. Estimates of the prevalence of BPD within the general population range from 0.5-1.5%^1,2^. Bipolar disorder is highly heritable, with siblings of probands at an 8-fold increased risk of the disorder^1,2^, and twin studies producing strikingly high estimates of heritability, around 89-93%^1,3,4^. More recently, genetic studies of BPD have indicated SNP heritability estimates of 17-23%^5^.

Bipolar disorder encompasses diagnostically distinct subtypes; bipolar disorder type I (BD-I), characterized by full manic episodes, and bipolar disorder type II (BD-II), which includes both hypomania and recurrent depressive episodes^1,6,7^. Individuals with diagnostic features of both bipolar disorder and schizophrenia may additionally be diagnosed with schizoaffective disorder (SAB)^7^. Recent studies have indicated that these diagnostic distinctions may be borne out genetically; for example, BD-I is significantly more heritable than BD-II^5,8^, and there are distinct differences between polygenic risk profiles of individuals with BD-I compared to BD-II^6,8^. These diagnostic and genetic heterogeneities within bipolar disorder contribute to the complexity in identifying genetic associations with bipolar disorder. Additional complications arise due to the complex polygenic nature of the disorder, and the high degree of overlap, both diagnostically and genetically, with other psychiatric disorders such as Schizophrenia and Major Depressive Disorder^9–11^.

Global collaborative efforts over the last decade have enabled large collections of samples from individuals with BPD. Genome-wide associations studies (GWAS) of these collections have identified multiple BPD-associated loci throughout the genome^6,12–25^, most recently 30 novel loci identified in the PGC-BD GWAS^5^. Despite these advances in locus discovery, little is understood about the pathogenesis of bipolar disorder. It is likely that, in line with other psychiatric disorders, larger sample sizes will be required in order to identify additional risk loci^26^. However, even elegantly designed and well-powered GWAS studies will not necessarily identify biological mechanisms contributing to disease, as large lists of genomic loci may be uninformative, and require careful dissection and downstream analyses to identify truly disease-causing associations^27^.

Transcriptomic Imputation (TI) analyses offer an opportunity to probe gene expression on a large scale, using eQTL reference panel-derived prediction models^28,29^. These approaches have several attractive advantages to researchers studying genetics of complex traits. First, results are readily biologically interpretable. Second, the large scale of GWAS studies means that TI studies are powered to detect even modest changes in gene expression, which likely represent a large portion of the risk in psychiatric disorders^30,31^, and which cannot be identified with traditional transcriptome approaches. Third, the use of genetically-regulated gene expression ensures that any associations precede symptom onset, rather than being mediated by disease status^28^.

In this study, we present the largest analysis of transcriptomic imputation in Bipolar Disorder. Our analysis included individuals from the most recent PGC-BD GWAS^5^ (19,986 cases/30,992 controls), as well as individuals from the iPSYCH consortium (1,502 cases/23,311 controls). We calculated predicted genetically regulated gene expression (GREX) for ~20,000 genes across 13 brain regions, using prediction models derived from GTEX^28,32^ and CommonMind Consortium data^31,33^. We sought to identify associations between GREX and a diagnosis of bipolar disorder, or one of three bipolar subtypes (BD-I, BD-II, SAB). We identified 125 significant gene-tissue associations with BPD, constituting 53 independent associations. Of these, 29 gene-tissue associations were novel; i.e., they did not lie within 1MB of a locus identified in the recent PGC-BD GWAS^5^. Additionally, we identified 80 gene-tissue associations with BD-I (37 independent associations, of which 12 were novel), two gene-tissue associations with BD-II (both novel), and one gene-tissue association with SAB. Our associations were highly consistent with differential gene expression analyses of bipolar cases and controls in the CommonMind Consortium. We expound upon these results using a number of analyses, including gene set enrichment analyses, replication of previous transcriptome-based studies of bipolar disorder^28,34^, and an approach analogous to PHEWAS^35,36^ to identify associations between these genes and specific endophenotypes of bipolar disorder.

## Methods

### Samples

Genotype data were obtained from the Psychiatric Genomics Consortium Bipolar Disorder (PGC-BD) collection. These data included 19,986 cases and 30,992 ancestry-matched controls from the PGC-BD collection^5^. Three of these cohorts were available through summary statistics only (Supplementary Figure 1). 1,502 BPD cases and 23,311 matched controls were additionally analysed by collaborators at iPSYCH (supplementary information).

In order to be included in the study, cases were required to meet international diagnostic criteria for BPD (ie, DSM-IV, ICD-9, ICD-10), or to have a lifetime diagnosis of BPD according to structured diagnostic instruments^5^. Genotyping information for these samples can be found in the flagship papers describing the initial sample collection^5^, and were processed in a standardized manner using “ricopili” ^5^.

The PGC-BD collection included 14,938 individuals with BD-I, 3,543 individuals with BD-II, and 1, 500 individuals with SAB. No subtype data were available for individuals collected through iPSYCH.

### Transcriptomic Imputation

We imputed genetically regulated gene expression (GREX) using the CommonMind Consortium (CMC) derived Dorso-lateral pre-frontal cortex (DLPFC) predictor model^33^, and GTEx-derived brain tissue prediction models^28,32^. We imputed GREX in all cohorts for which we had access to raw data using PrediXcan^28^ (Suppl. Figure 1).

For three cohorts, raw genotype data was not available. For these cohorts, and two cohorts with a trio structure, genic associations were computed using summary statistics, using MetaXcan^37^, a summary-statistic approach analogous to prediXcan^28^. Previous studies have shown that genic association p-values and effect sizes calculated using MetaXcan and PrediXcan are highly correlated, provided that ethnically matched reference panels are used^33,37^. This was confirmed using three European PGC BD cohorts for which both summary statistics and raw genotype data were available.

### iPsych-Gems Analysis

iPSYCH-GEMS GWAS data was genotyped and imputed in 23 waves, and subsequently merged for association analyses. No subtype data were available for iPSYCH-GEMS data. Variants with imputation scores>0.8 were included for the analysis. Genetically regulated gene expression levels were calculated using the CMC DLPFC predictor model^33^, as well as 12 GTEx-derived brain tissue databases^28,32^. Association tests on case-control status were carried out using a logistic regression in R, including wave membership as covariate.

Principal component analysis was done in order to remove genetic outliers. The phenotype specific PCs that are significantly different between cases and controls were included as covariates as well, to account for the population stratification. Related individuals were identified by pairwise IBD analysis and one of every pair (preferably controls) identified as related (piHAT > 0.2) was removed.

Regression formula: Disease ~ gene-expression + wave1 + wave2 + …. + wave22 + PC1+PC2+…

The association analysis was done using R software.

### Association Tests

We tested for association between GREX and case-control status in each cohort separately, using a standard linear regression test in R. We included ten principal components as covariates. We repeated this analysis for BD-I, BD-II and SAB, including all controls. We required that a cohort include at least 50 individuals with a given subtype to be included in each analysis, and consequently removed one cohort with only 36 SAB cases.

We carried out an analysis comparing bipolar subtypes BD-I, BD-II, SAB. For each pair of subtypes, we compared GREX in cases only, including all cohorts with more than 50 individuals with each diagnosis.

Raw genotype-based and summary-statistics based cohorts were meta-analysed using an odds-ratio based approach in METAL^38^.

### Establishing a threshold for genome-wide significance

We applied two significance thresholds to the data. First, for each tissue, we applied a Bonferroni correction accounting for the total number of genes tested within that tissue (Suppl. table 1). Second, we applied a global genome-wide significance threshold, accounting for all genes tested across all tissues. These are denoted by dashed and solid lines respectively in the manhattan plots throughout this manuscript.

### Identifying independent associations

We identified 18 regions with multiple gene-tissue associations; regions were defined based on distance between genes, and were checked using visual inspection of associations across each chromosome. For each of these regions, we applied FINEMAP^39^ to identify independently associated genes. We substituted the LD-matrix usually used in FINEMAP with an analogous GREX correlation matrix.

This matrix was calculated for each cohort with available raw data, and a weighted average calculated across all populations, weighting for effective sample size. We ensured that summary-statistic based cohorts were represented in this weighted average by selecting the geographically nearest cohort as a proxy, and increasing the weighting of that proxy cohort accordingly.

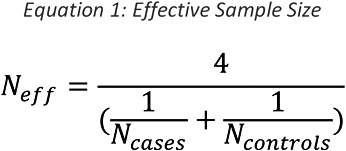

### Identifying genes associated with specific behaviours and clinical variables

We obtained data on 26 clinical variables relating to BPD, including for example rapid cycling, psychosis, panic attacks, and a variety of comorbid disorders. We used an approach analogous to PHEWAS, and an adaptation to the PHEWAS R package^40^, to test for associations between BD-I, BD-II and SAB-associated genes and these 26 endophenotypes.

Behavioural data was available for ~8,500 individuals, across 14 cohorts. We tested for association between GREX and all 26 endophenotypes in each cohort separately, controlling for ten principal components. Only endophenotypes with at least 20 cases, or 20 quantitative measures, were included within each cohort. Results were meta-analyzed across cohorts using an odds-ratio based approach in METAL^41^.

### Comparison with Differential Expression in CommonMind Consortium

We sought to compare putatively BPD-associated GREX changes to genes identified as differentially expressed in post-mortem brain samples. We obtained summary statistics on differential expression between Bipolar cases and healthy controls from the CommonMind Consortium Phase II analysis, across the dorso-lateral pre-frontal cortex (DLPFC; 55 cases, 296 controls) and anterior cingulate cortex (ACC; 48 cases, 246 controls).

We compared association statistics between these two analyses and each of our prediXcan BPD analyses; specifically, we tested whether genes reaching tissue-specific significance in each prediXcan analysis were more likely than expected by chance to be differentially expressed in the CMC analysis. We then repeated this test using all nominally significant genes in the prediXcan analyses. Additionally, we tested whether the degree of replication seen in each tissue was correlated with the number of genes tested, and/or with the sample size of the original eQTL reference panel used.

Since we did not have access to individual-level RNA-seq data in order to run a BD-I specific differential expression analysis, we compared BD-I DLPFC and ACC prediXcan association statistics to the CMC differential expression analysis.

We identified a small number of individuals within the CommonMind Consortium sample who were diagnosed with BD-II subtype. No RNA-seq data was available for these individuals; however, 11 had available microarray data. We therefore compared normalized microarray data between these 11 individuals and 204 controls, for the two top genes in our BD-II subtype analysis *(COLGALT2* and *NUP98).* No individuals with SAB were available for analysis.

### Pathway Analysis

Pathway analysis was carried out using an adaptation to MAGMA^42^. We performed three pathway analyses, as follows: 1) 174 drug-target gene sets; 2) 76 gene sets with prior evidence of involvement in BD^31,43–45^, including nervous-systems related pathways, gene sets relating to aberrant behavior in mice, circadian clock gene sets, calcium-gated voltage channels, as well as targets of FMRP; 3) ~8,500 pathways collated across six large publicly available datasets^46–53^. We included only gene sets with at least 10 genes.

For each of the four iterations, we analyzed BIP, BD-I, BD-II and SAB results separately. Analyses were carried out using genic p-values from our PrediXcan meta-analyses. In instances where a gene had multiple associations across different tissues, the best p-value was selected, and a Bonferroni correction applied to correct for the number of tissues tested. Gene-set enrichment results from the competitive (rather than self-contained) MAGMA analysis were used^42^, and FDR correction applied within each stratum of our analysis.

## Results

### Association Tests

We calculated predicted gene expression for thirteen brain regions (derived from CMC and GTEx data^28,32,54,55^) in 19,986 cases and 30,992 controls from the PGC-BPD^5^ and 1,502 cases and 23,311 controls from the iPsych-GEMS consortium, and tested for association between predicted gene expression (GREX) and case-control status. Additionally, we used a summary-statistic based method to calculate genic associations in cases and controls for which raw genotypes were not available (Suppl. Figure 1A).

We identified 125 genes-tissue associations reaching tissue-specific significance (Suppl. Table 2; Figure 1A; ~5e-06); 46/125 reached our stricter cross-tissue threshold (4.11e-07). Within these associations, we identified 18 genomic regions with multiple associated genes, and where the same gene was associated across multiple tissues. We applied FINEMAP to each of these regions, and identified 53 independent associations (Table 1; Figure 1B), of which 29 are novel (i.e., they do not lie within 1Mb of a locus identified in the recent PGC-BD GWAS^5^). It should be noted that our sample includes all of the PGC-BD samples as well as an additional cohort, and so will have greater power to detect signals than the original GWAS.

**Figure 1:**
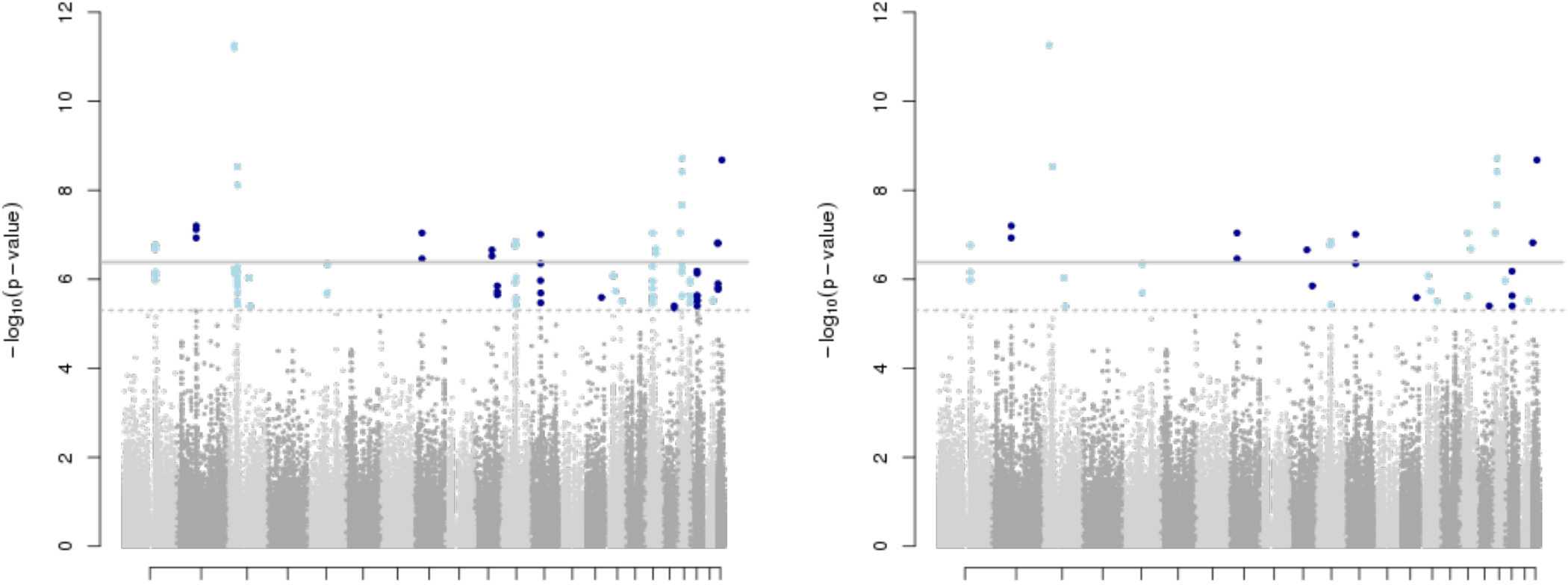
Genic associations identified across full Bipolar sample. A) 125 gene-tissue associations are identified in the full BPD meta-analysis B) FINEMAP analysis identifies 53 independent associations

**Table 1:**
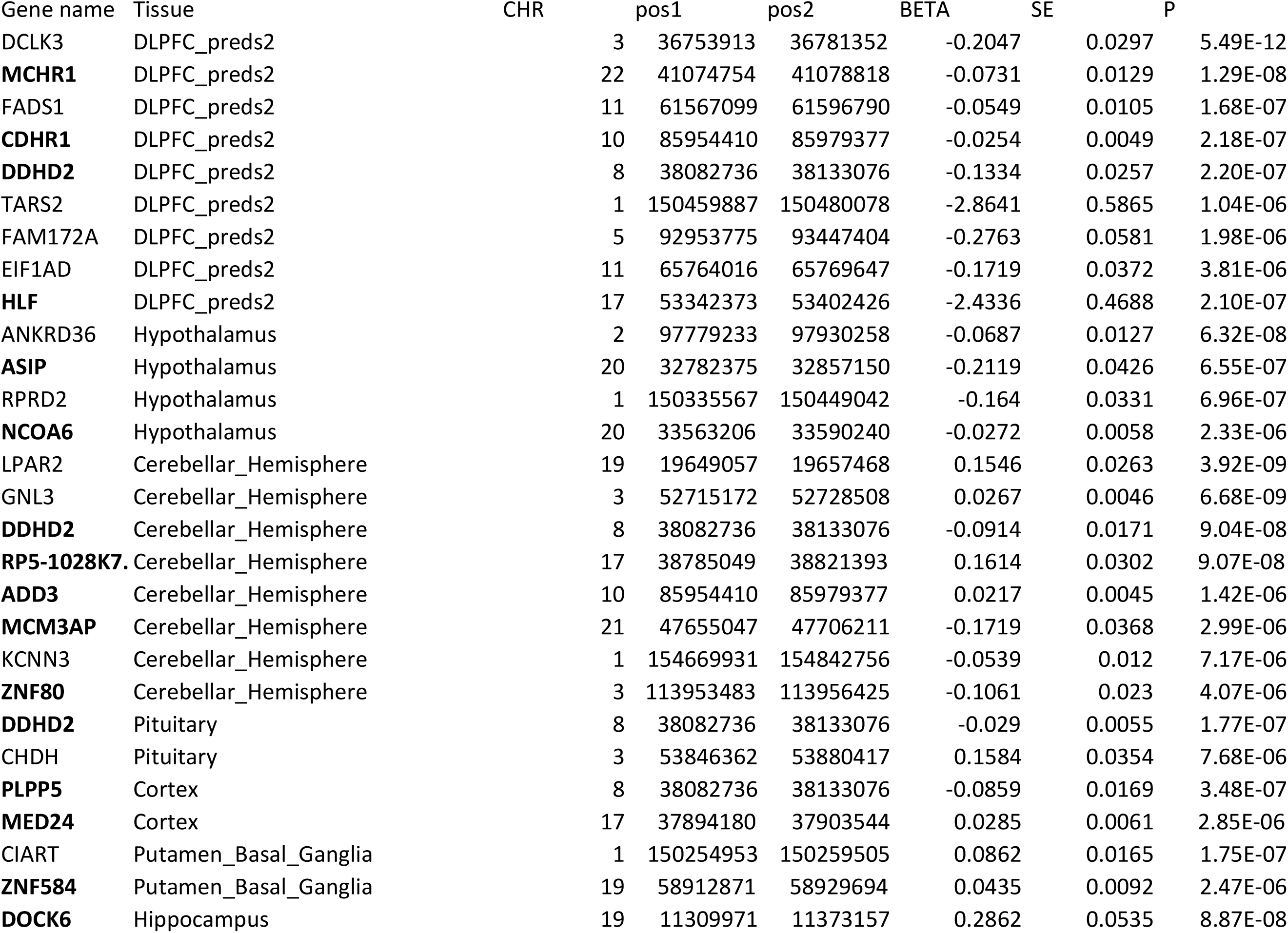
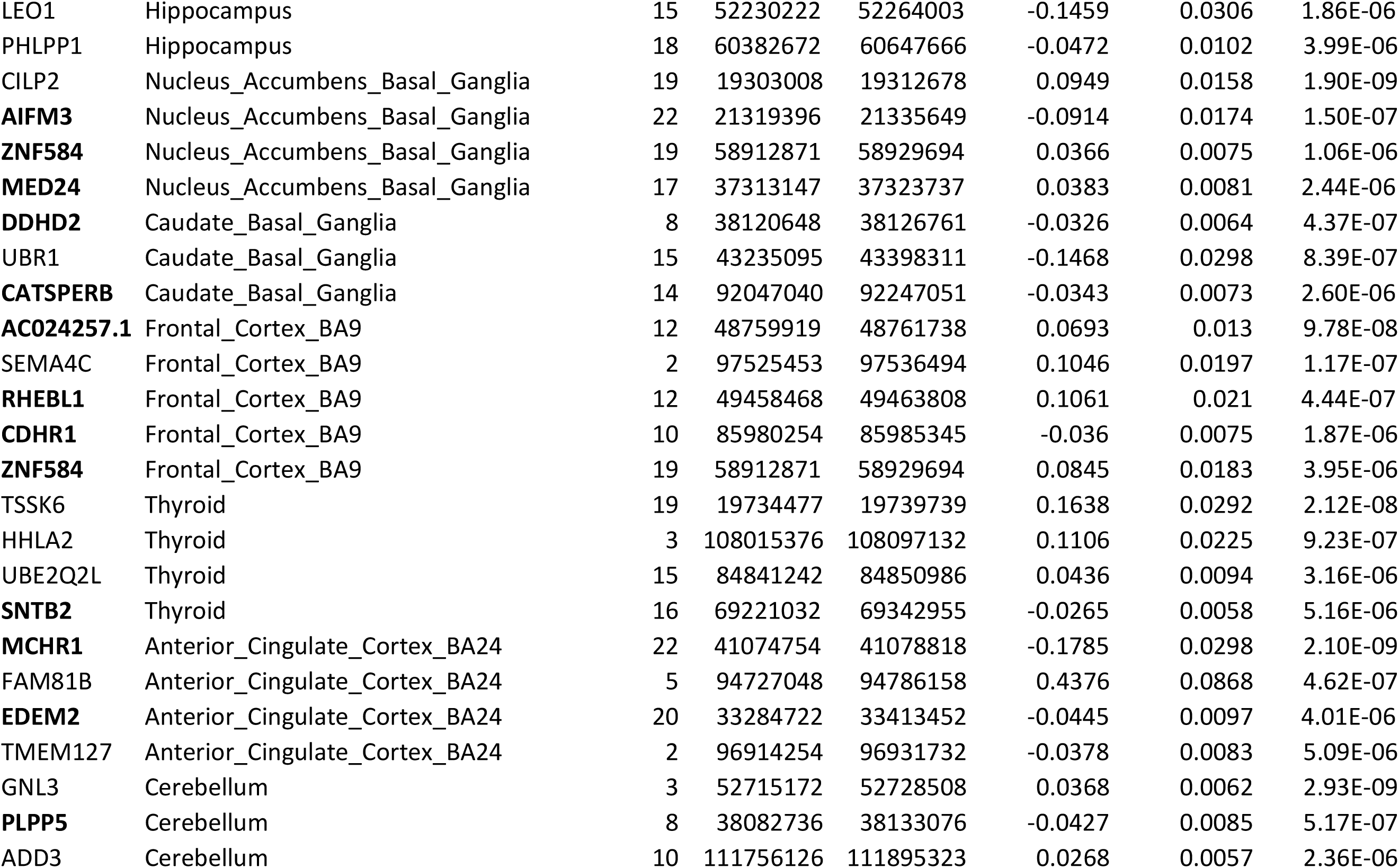
Gene-Tissue Associations results

### Comparison to previous transcriptome studies

Two previous studies have already identified BPD-associated genes using transcriptomic approaches, albeit using substantially smaller samples^28,34^. We sought to replicate these findings using the subset of our data not included in the original PGC-BD GWAS^5^ (Table 2).

**Table 2:** *Replication p-values of genes identified in previous Transcriptome Analysis of BPD*

| Gene | Tissue | p-value | Direction of Effect |
| --- | --- | --- | --- |
| PTPRE | Putamen Basal Ganglia | 0.024 | - |
| SPCS1 | **Caudate Basal Ganglia** | **0.0011** | **+** |
| CACNB3 | **Frontal Cortex BA9** | **0.0010** | - |
|  | Anterior Cingulate Cortex | 0.0032 | - |
|  | Whole Blood | 0.0042 | + |
|  | Cerebellum | 0.0044 | - |
|  | Cerebellar Hemisphere | 0.0080 | - |
|  | Caudate Basal Ganglia | 0.012 | - |
|  | DLPFC | 0.019 | - |
|  | Nucleus Accumbens Basal Ganglia | 0.027 | - |
|  | Putamen Basal Ganglia | 0.077 | - |

One gene, *PTPRE*, was identified as associated with Bipolar Disorder in the original prediXcan-based Transcriptomic Imputation analysis. Two genes, *SPCS1* and *CACNB3*, were identified using the SMR method^34^, which used eQTLs from peripheral blood. *PTPRE* reaches nominal significance in the putamen basal ganglia in our replication sample (p=0.024). Both *SPCS1* and *CACNB3* were significant in our replication sample (after Bonferroni correction); *SPCS1* in the caudate basal ganglia (p=0.0011), and *CACNB3* in the frontal cortex (p=0.0010). Additionally, *CACNB3* reaches nominal significance in seven other tissues. This level of replication is highly unlikely to occur by chance (binomial test: p=1.59x10^−7^ at nominal significance threshold, p=0.0012 at Bonferroni-corrected threshold).

### Subtypes

Bipolar disorder subtypes BD-I, BD-II and SAB have previously been shown to be diagnostically and genetically distinct^6^. We tested for association of GREX with case-control status for each of these three subtypes, using all available matched controls; BD-I (14,983 cases/controls), BD-II (3,543/22,155) and SAB (1,500/8,690).

We identified 80 BD-I gene-tissue associations reaching tissue-specific genome-wide significance (~6x10^−06^; Suppl. Table 3), constituting 37 independent associations following FINEMAP (Table 3; Figure 2A). 12 gene-tissue associations across 10 regions were novel, i.e., did not lie within 1Mb of a BD-I locus identified in the PGC-BD GWAS^5^. In line with our overall BPD analysis, the largest number of associations occur in the cortex and pre-frontal cortex (14 associations) and the limbic system (14 associations).

**Figure 2:**
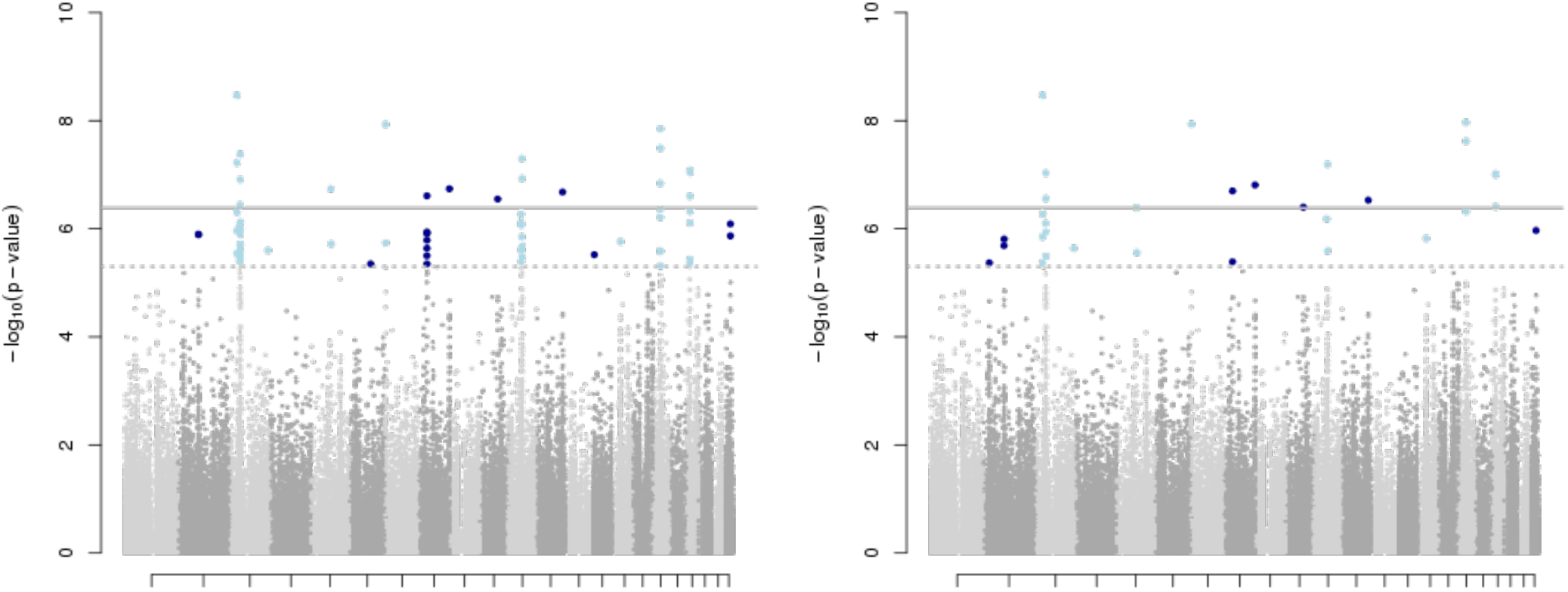
Genic associations identified in three bipolar subtypes. A) 80 gene-tissue associations are identified in the Bipolar-I sample. B) FINEMAP and Stepwise conditional analysis identify 37independent associations

**Table 3:**
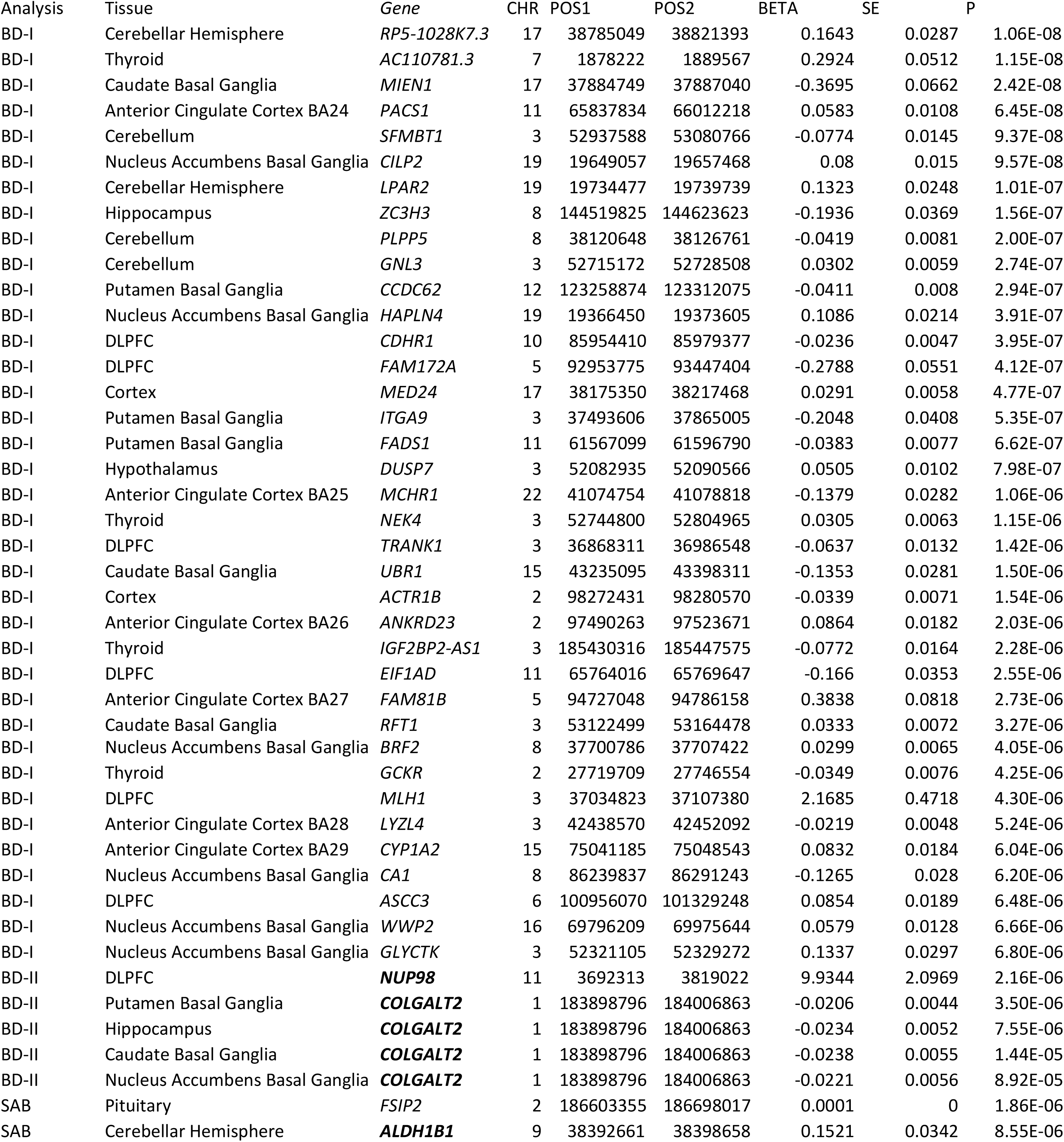
Gene-Tissue Associations results for subtype analyses

Two genes were associated with BD-II subtype, albeit not at the stricter cross-tissue significance threshold (Table 3). First, increased expression *NUP98* in the DLPFC was associated with BD-II (p=2.2e-06). Decreased expression of *COLGALT2* was associated with BD-II in the Putamen Basal Ganglia (p=3.5e-06) and neared significance in the Hippocampus (p=7.6e-06), the Caudate Basal Ganglia (p=1.4e-05) and the Nucleus Accumbens Basal Ganglia (p=8.9e-05). Neither of these BD-II genes lie within 1Mb of a BD-II locus identified in the recent PGC-BD GWAS, although other BD-II subthreshold associations do (Suppl. Table 4).

Increased expression of *FSIP2* in the Thyroid was associated with SAB (p=1.9e-06; Table 3). Increased expression of *ALDH1B1* in the Cerebellar Hemisphere was also associated with SAB, although at slightly below tissue-specific significance (p=8.4e-06). *FSIP2* lies ~0.5Mb from a locus also identified as potentially associated with SAB in the PGC-BD GWAS (p=6.9x10^−7^). One sub-threshold association *(SNX29*, in the Hypothalamus; Suppl. Table 4), also lies close to a PGC-BD GWAS SAB locus; all other SAB associations are novel.

There is a substantial overlap between association signals in our BD and BD-I analyses, likely due to the high proportion of BD-I cases within our sample, and a high proportion of overlapping controls. We examined association statistics (-log10 p-values) of all associated genes across all four analyses (Figure 3) and noted that BD and BD-1 genes tend to be reciprocally associated, whereas genes identified in the BD-2 and SAB analyses tend to be associated only within those particular subtypes.

**Figure 3:**
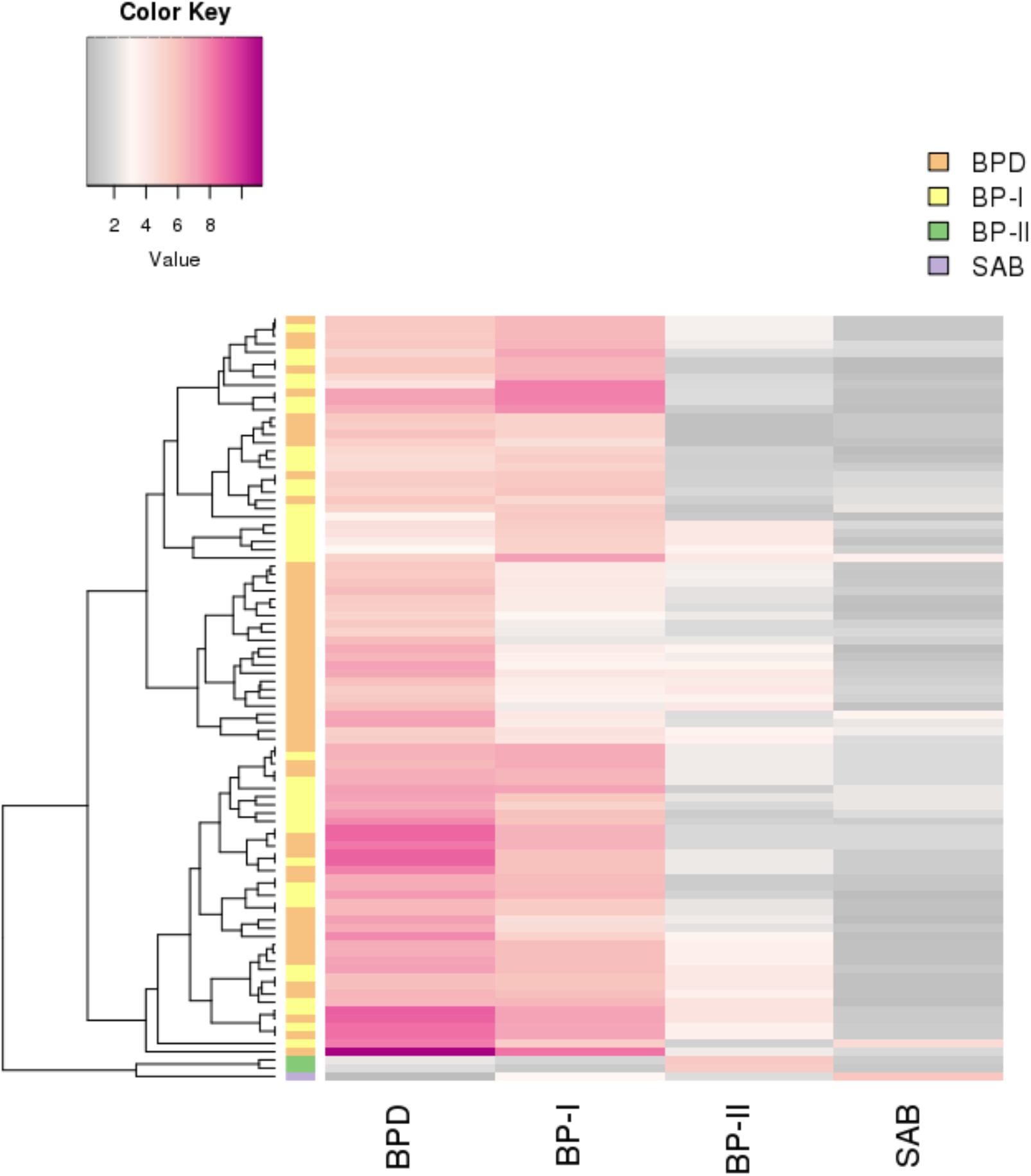
Substantial overlap between BPD and BP-I associated genes. -logl0 p-values are shown for all genes reaching genome-wide significance in any discovery analysis. The row side colour bar indicates the original discovery analysis identifying the gene. The four row values indicate the best p-value achieved by that gene in each subtype analysis. e.g.: the bottom row shows a gene *(FSIP2)* identified in the SAB subtype analysis, and the best p-value achieved by *FSIP2* across all tissues in the overall BPD analysis, BD-I, BD-II and SAB analyses.

### Comparison to Differential Expression in the CommonMind Consortium samples

We compared our prediXcan GREX results to bipolar disorder differential expression analysis conducted in CommonMind Consortium post-mortem samples. Across all tissues, genes reaching nominal significance in our prediXcan analysis were significantly more likely to be differentially expressed in CMC DLPFC post-mortem samples (binomial test, p<2.8e-73; Supplementary Table 5). The degree of replication was significantly correlated with the sample size of the original eQTL reference panel, even when controlling for the number of genes tested (p=0.03).

Genes reaching tissue-specific significance (p<0.05/N genes tested) in the DLPFC, ACC, Cortex, and Nucleus Accumbens prediXcan analyses were more likely than expected by chance to be differentially expressed in the DLPFC CMC post-mortem samples (binomial test, p<0.0038). There was no relationship between the likelihood of replication of significant genes and the number of genes tested, or eQTL reference panel sample size.

The vast majority of BPD cases in the CommonMind Consortium differential expression analysis were BD-I subtype; therefore, we also used the same CMC differential expression analysis to test for replication of our BD-I prediXcan results. As for the overall BPD analysis, nominally significant prediXcan genes were all significantly more likely to be differentially expressed in our CMC analysis (binomial test, p<4.57e-72), and the degree of replication was correlated with sample size of the original eQTL reference panel (p=0.044). Genes reaching tissue-specific significance in both the DLPFC and the Cortex were significantly more likely to be differentially expressed in the CMC analysis (binomial test, p<0.0016; Supplementary Table 5).

We identified a small number of individuals within the CommonMind Consortium sample who were diagnosed with BD-II subtype. No RNA-seq data was available for these individuals; however, 11 had available microarray expression data. We therefore compared normalized microarray data between these 11 individuals and 204 controls, for the two top genes in our BD-II subtype analysis *(COLGALT2* and *NUP98).* Both genes had the same directions of effect between cases and controls in our CMC Microarray data as in the prediXcan meta-analysis. In particular, the ratio of case:control expression for *COLGALT2* was strikingly similar in the microarray data (0.984) to the effect size estimated using prediXcan (0.980), and expression levels were significantly different between cases and controls (p=0.0488). However, the sample sizes in this analysis are small, and results should be taken as preliminary, exploratory findings, and further, larger analysis will be required.

No individuals with SAB were available for analysis.

### Identifying genes associated with specific behaviours

We tested whether any of the genes identified in our subtype analyses were particularly associated with any specific BPD-endophenotype, using an approach analogous to PHEWAS^35,36^. We included all genes reaching tissue-specific significance in any subtype analysis.

We identified three significant associations (Table 4). We found that reduced expression of *EIF1AD* in the DLPFC was associated with mixed states (p=0.00197) and panic attacks (p=0.0004948). In our original analysis, decreased expression of the gene in the DLPFC was associated with BD-I (p=2.55x10^−6^). Additionally, decreased expression of *FSIP2* in the Pituitary was associated with having a family history of BPD in our PHEWAS (p=1e-05).

**Table 4:** Endophenotype-wide association study (enPHEWAS). All genes reaching tissue-wide significance in any subphenotype-based analysis were included.

| Gene | Tissue | enPHEWAS Analysis |  |  |  |  | Subtype-specific meta-analysis |  |  |  |  |
| --- | --- | --- | --- | --- | --- | --- | --- | --- | --- | --- | --- |
|  |  | Endophenotype | beta | se | p | OR | Subtype | beta | se | p | OR |
| *EIF1AD* | DLPFC | mixedstates | -0.3873 | 0.1252 | 1.97E-03 | 0.68 | BD-I | -0.166 | 0.0353 | 2.55E-06 | 0.85 |
| *EIF1AD* | DLPFC | panic.attacks | -0.2861 | 0.0821 | 4.95E-04 | 0.75 | BD-I | -0.166 | 0.0353 | 2.55E-06 | 0.85 |
| *FAM172A* | DLPFC | bp2 | 0.127 | 0.0393 | 1.24E-03 | 1.14 | BD-I | -0.2788 | 0.0551 | 4.12E-07 | 0.76 |
| *FSIP2* | Pituitary | *famhistory* | -0.0009 | 0.0002 | 1.09E-05 | 1.00 | SAB | 0.0001 | 0 | 1.86E-06 | 1.00 |

### Pathway enrichment

We tested for pathway enrichment using MAGMA^42^, for BD, BD-I, BD-II and SAB associations. We carried out three stages of pathway analysis including the following gene sets 1) 174 sets of drug targets; 2) 79 hypothesis-driven gene sets including targets of the FMRP protein, calcium-gated voltage channels, pathways involved in aberrant mouse behavior, pathways pertaining to chronotype and circadian rhythms 3) ~8,500 agnostic pathways obtained from large publicly available databases. All FDR-corrected significant results for these analyses are shown in Table 5.

**Table 5:** Pathway Results

| Association statistics | Analysis type | SET | NGENES | COMP P | FDR |
| --- | --- | --- | --- | --- | --- |
| BPD | Drug targets | ANABOLIC STEROIDS | 34 | 4.02E-06 | 0.001 |
| BPD | Drug targets | CORTICOSTEROIDS FOR SYSTEMIC USE PLAIN | 43 | 8.84E-05 | 0.013 |
| BPD | Drug targets | ANDROGENS | 47 | 1.72E-04 | 0.025 |
| BPD | Drug targets | ANTIFUNGALS FOR TOPICAL USE | 92 | 4.48E-04 | 0.064 |
| BPD | Hypothesis driven | MORNING | 109 | 3.27E-05 | 0.003 |
| BPD | Hypothesis driven | HIGH | 2718 | 1.08E-03 | 0.029 |
| BPD | Hypothesis driven | SCZ-NS | 567 | 1.29E-03 | 0.029 |
| BPD | Hypothesis driven | FMRP-targets | 735 | 1.47E-03 | 0.029 |
| BPD | Hypothesis driven | Pre-synaptic active zone | 156 | 4.20E-03 | 0.066 |
| BPD | Hypothesis driven | Circadian clock genes | 380 | 1.21E-02 | 0.159 |
| BPD | Hypothesis driven | CLOCK-MODULATORS | 254 | 2.32E-02 | 0.262 |
| BPD | Hypothesis driven | PSD-95 (core) | 56 | 3.52E-02 | 0.348 |
| BPD | Hypothesis driven | ID-LoF | 26 | 4.34E-02 | 0.381 |
| BPD | Hypothesis driven | ARC+NMDAR+PSD95+mGluR5 | 122 | 5.45E-02 | 0.416 |
| BPD | Hypothesis driven | SCZ-LoF | 79 | 5.79E-02 | 0.416 |
| BPD | Hypothesis driven | Cav2::kinases & phosph... | 20 | 8.99E-02 | 0.504 |
| BPD | Hypothesis driven | ID-NS | 116 | 9.92E-02 | 0.504 |
| BD-I | Agnostic | S-adenosylmethionine-dependent methyltransfe | 91 | 3.76E-08 | 0.000 |
| BD-I | Agnostic | mitochondrial nucleoid | 33 | 5.64E-07 | 0.001 |
| BD-I | Agnostic | nucleoid | 34 | 8.11E-07 | 0.001 |
| BD-I | Agnostic | RNA methylation | 27 | 9.35E-07 | 0.001 |
| BD-I | Agnostic | N-methyltransferase activity | 59 | 9.64E-07 | 0.001 |
| BD-I | Agnostic | RNA methyltransferase activity | 26 | 9.73E-07 | 0.001 |
| BD-I | Agnostic | regulation of transcription from RNA polymerase | 16 | 5.25E-06 | 0.006 |
| BD-I | Agnostic | impaired wound healing | 25 | 8.91E-06 | 0.010 |
| BD-I | Agnostic | Downregulation of ERBB2:ERBB3 signaling | 13 | 2.53E-05 | 0.020 |
| BD-I | Agnostic | extracellular regulation of signal transduction | 15 | 2.55E-05 | 0.020 |
| BD-I | Agnostic | extracellular negative regulation of signal transd | 15 | 2.55E-05 | 0.020 |
| BD-I | Agnostic | abnormal cellular respiration | 66 | 2.92E-05 | 0.021 |
| BD-I | Agnostic | male meiosis | 32 | 3.91E-05 | 0.026 |
| BD-I | Agnostic | Fanconi Anemia pathway | 21 | 5.83E-05 | 0.036 |
| BD-I | Agnostic | Golgi-associated vesicle | 56 | 7.65E-05 | 0.043 |
| BD-I | Agnostic | regulation of T cell migration | 13 | 8.00E-05 | 0.043 |
| BD-I | Agnostic | positive regulation of T cell migration | 11 | 1.01E-04 | 0.051 |
| BD-I | Agnostic | failure of tooth eruption | 16 | 1.34E-04 | 0.064 |
| BD-I | Agnostic | viral assembly | 29 | 1.43E-04 | 0.064 |
| BD-I | Agnostic | macromolecule methylation | 138 | 1.61E-04 | 0.066 |
| BD-I | Agnostic | viral infectious cycle | 121 | 1.67E-04 | 0.066 |
| BD-I | Agnostic | negative regulation by host of viral transcription | 12 | 1.69E-04 | 0.066 |
| BD-I | Agnostic | skeletal muscle contraction | 16 | 1.93E-04 | 0.072 |
| BD-I | Agnostic | toxin metabolic process | 10 | 2.77E-04 | 0.096 |
| BD-I | Agnostic | granulomatous inflammation | 24 | 2.79E-04 | 0.096 |
| BD-I | Hypothesis driven | Endoplasmic Reticulum (core) | 87 | 2.04E-04 | 0.036 |
| BD-I | Hypothesis driven | PSD (human core) | 624 | 5.16E-04 | 0.046 |
| BD-I | Hypothesis driven | MORNING | 109 | 1.16E-03 | 0.069 |
| BD-II | Agnostic | S-adenosylmethionine-dependent methyltransfe | 91 | 3.46E-07 | 0.003 |
| BD-II | Agnostic | negative regulation of systemic arterial blood pr | 11 | 2.09E-06 | 0.009 |
| BD-II | Agnostic | Metabolism of porphyrins | 15 | 4.92E-06 | 0.014 |
| BD-II | Agnostic | RNA polymerase activity | 37 | 9.47E-06 | 0.016 |
| BD-II | Agnostic | DNA-directed RNA polymerase activity | 37 | 9.47E-06 | 0.016 |
| BD-II | Agnostic | Endogenous sterols | 15 | 1.83E-05 | 0.026 |
| BD-II | Agnostic | Heme biosynthesis | 12 | 2.38E-05 | 0.027 |
| BD-II | Agnostic | protein methyltransferase activity | 58 | 2.54E-05 | 0.027 |
| BD-II | Agnostic | condensed chromosome | 145 | 4.07E-05 | 0.032 |
| BD-II | Agnostic | mitochondrial genome maintenance | 12 | 4.08E-05 | 0.032 |
| BD-II | Agnostic | nuclear envelope organization | 52 | 4.46E-05 | 0.032 |
| BD-II | Agnostic | centrosome localization | 12 | 5.04E-05 | 0.032 |
| BD-II | Agnostic | abnormal nucleotide metabolism | 10 | 5.28E-05 | 0.032 |
| BD-II | Agnostic | chondroitin sulfate metabolic process | 50 | 5.28E-05 | 0.032 |
| BD-II | Agnostic | heme metabolic process | 29 | 6.75E-05 | 0.038 |
| BD-II | Agnostic | porphyrin-containing compound metabolic process | 38 | 7.74E-05 | 0.038 |
| BD-II | Agnostic | protoporphyrinogen IX metabolic process | 10 | 7.83E-05 | 0.038 |
| BD-II | Agnostic | chondroitin sulfate biosynthetic process | 23 | 9.52E-05 | 0.038 |
| BD-II | Agnostic | abnormal spinal cord morphology | 193 | 9.60E-05 | 0.038 |
| BD-II | Agnostic | Heme biosynthesis | 10 | 9.78E-05 | 0.038 |
| BD-II | Agnostic | N-methyltransferase activity | 59 | 1.01E-04 | 0.038 |
| BD-II | Agnostic | chondroitin sulfate proteoglycan metabolic process | 52 | 1.07E-04 | 0.038 |
| BD-II | Agnostic | abnormal neuronal migration | 75 | 1.09E-04 | 0.038 |
| BD-II | Agnostic | nucleotidyltransferase activity | 106 | 1.10E-04 | 0.038 |
| BD-II | Agnostic | Chondroitin sulfate | 43 | 1.12E-04 | 0.038 |
| BD-II | Agnostic | nucleoside kinase activity | 11 | 1.14E-04 | 0.038 |
| BD-II | Agnostic | KEGG PYRIMIDINE METABOLISM | 90 | 1.50E-04 | 0.048 |
| BD-II | Agnostic | oxidoreductase activity | 11 | 1.56E-04 | 0.048 |
| BD-II | Agnostic | Transport of Mature mRNA Derived from an Intronic | 34 | 1.94E-04 | 0.057 |
| BD-II | Agnostic | nucleoside salvage | 12 | 2.06E-04 | 0.059 |
| BD-II | Agnostic | neuron spine | 69 | 2.47E-04 | 0.068 |
| BD-II | Agnostic | protein methylation | 82 | 2.71E-04 | 0.070 |
| BD-II | Agnostic | protein alkylation | 82 | 2.71E-04 | 0.070 |
| BD-II | Agnostic | dendritic spine | 67 | 2.83E-04 | 0.071 |
| BD-II | Agnostic | chondroitin sulfate proteoglycan biosynthetic process | 26 | 2.95E-04 | 0.072 |
| BD-II | Agnostic | RNA splicing | 179 | 3.66E-04 | 0.085 |
| BD-II | Agnostic | mRNA splicing | 179 | 3.66E-04 | 0.085 |
| BD-II | Agnostic | abnormal mitochondrion morphology | 55 | 3.89E-04 | 0.088 |
| BD-II | Drug targets | THYROID PREPARATIONS | 11 | 1.65E-09 | 0.000 |
| SAB | Agnostic | laminin complex | 10 | 6.64E-08 | 0.001 |
| SAB | Agnostic | abnormal miniature endplate potential | 21 | 1.54E-07 | 0.001 |
| SAB | Agnostic | a6b1 a6b4 integrin pathway | 46 | 1.78E-07 | 0.001 |
| SAB | Agnostic | Schwann cell development | 20 | 2.59E-07 | 0.001 |
| SAB | Agnostic | astrocyte development | 12 | 3.19E-07 | 0.001 |
| SAB | Agnostic | abnormal amacrine cell number | 11 | 4.32E-07 | 0.001 |
| SAB | Agnostic | glomerular basement membrane development | 10 | 8.59E-07 | 0.001 |
| SAB | Agnostic | cellular amide metabolic process | 138 | 1.05E-06 | 0.001 |
| SAB | Agnostic | abnormal PNS synaptic transmission | 28 | 1.92E-06 | 0.002 |
| SAB | Agnostic | short photoreceptor inner segment | 13 | 2.74E-06 | 0.002 |
| SAB | Agnostic | renal filtration cell differentiation | 13 | 3.54E-06 | 0.003 |
| SAB | Agnostic | glomerular visceral epithelial cell differentiation | 13 | 3.54E-06 | 0.003 |
| SAB | Agnostic | abnormal photoreceptor inner segment morpho | 33 | 4.01E-06 | 0.003 |
| SAB | Agnostic | Schwann cell differentiation | 26 | 4.70E-06 | 0.003 |
| SAB | Agnostic | axon regeneration | 15 | 4.71E-06 | 0.003 |
| SAB | Agnostic | abnormal Muller cell morphology | 10 | 5.16E-06 | 0.003 |
| SAB | Agnostic | glomerular epithelium development | 14 | 6.23E-06 | 0.003 |
| SAB | Agnostic | glomerular epithelial cell differentiation | 14 | 6.23E-06 | 0.003 |
| SAB | Agnostic | myofilament | 17 | 9.13E-06 | 0.004 |
| SAB | Agnostic | striated muscle thin filament | 14 | 1.28E-05 | 0.005 |
| SAB | Agnostic | mitochondrion localization | 17 | 2.53E-05 | 0.010 |
| SAB | Agnostic | abnormal retinal apoptosis | 31 | 2.79E-05 | 0.011 |
| SAB | Agnostic | neuron projection regeneration | 21 | 3.25E-05 | 0.012 |
| SAB | Agnostic | abnormal amacrine cell morphology | 19 | 3.37E-05 | 0.012 |
| SAB | Agnostic | neuromuscular junction development | 33 | 5.04E-05 | 0.017 |
| SAB | Agnostic | integral to lumenal side of endoplasmic reticular | 24 | 5.87E-05 | 0.019 |
| SAB | Agnostic | endoplasmic reticulum-Golgi intermediate comp | 24 | 8.57E-05 | 0.027 |
| SAB | Agnostic | negative regulation of ion transmembrane trans | 11 | 9.33E-05 | 0.029 |
| SAB | Agnostic | biotin metabolic process | 10 | 1.06E-04 | 0.030 |
| SAB | Agnostic | Biotin transport and metabolism | 10 | 1.06E-04 | 0.030 |
| SAB | Agnostic | Methylation | 10 | 1.18E-04 | 0.033 |
| SAB | Agnostic | abnormal retinal rod cell morphology | 36 | 1.30E-04 | 0.035 |
| SAB | Agnostic | cell differentiation involved in metanephros dev | 11 | 1.33E-04 | 0.035 |
| SAB | Agnostic | secondary metabolic process | 68 | 1.52E-04 | 0.038 |
| SAB | Agnostic | branched-chain amino acid catabolic process | 18 | 1.76E-04 | 0.042 |
| SAB | Agnostic | negative regulation of transmembrane transport | 13 | 1.78E-04 | 0.042 |
| SAB | Agnostic | metanephric glomerulus development | 10 | 2.27E-04 | 0.052 |
| SAB | Agnostic | regulation of DNA-dependent transcription in re: | 41 | 2.32E-04 | 0.052 |
| SAB | Agnostic | branched-chain amino acid metabolic process | 22 | 2.42E-04 | 0.053 |
| SAB | Agnostic | tropomyosin binding | 14 | 2.74E-04 | 0.058 |
| SAB | Agnostic | positive regulation of receptor biosynthetic proc | 10 | 2.85E-04 | 0.058 |
| SAB | Agnostic | cellular amino acid biosynthetic process | 97 | 2.95E-04 | 0.058 |
| SAB | Agnostic | skeletal muscle fiber development | 44 | 3.04E-04 | 0.058 |
| SAB | Agnostic | Branched-chain amino acid catabolism | 16 | 3.07E-04 | 0.058 |
| SAB | Agnostic | KEGG TOXOPLASMOSIS | 110 | 3.07E-04 | 0.058 |
| SAB | Agnostic | negative regulation of GTPase activity | 15 | 3.29E-04 | 0.061 |
| SAB | Agnostic | BIOCARTA ERK5 PATHWAY | 17 | 3.35E-04 | 0.061 |
| SAB | Agnostic | decreased cellular sensitivity to gamma-irradiati | 18 | 3.48E-04 | 0.061 |
| SAB | Agnostic | astrocyte differentiation | 23 | 3.50E-04 | 0.061 |
| SAB | Agnostic | positive regulation of glucose import | 27 | 3.82E-04 | 0.065 |
| SAB | Agnostic | photoreceptor connecting cilium | 21 | 3.96E-04 | 0.066 |
| SAB | Agnostic | short photoreceptor outer segment | 27 | 4.03E-04 | 0.066 |
| SAB | Agnostic | 14-3-3 protein binding | 16 | 4.51E-04 | 0.073 |
| SAB | Agnostic | protein phosphatase 2A binding | 16 | 4.98E-04 | 0.079 |
| SAB | Agnostic | fatty acid derivative metabolic process | 74 | 5.28E-04 | 0.081 |
| SAB | Agnostic | icosanoid metabolic process | 74 | 5.28E-04 | 0.081 |
| SAB | Agnostic | KEGG GLYCOSYLPHOSPHATIDYLINOSITOL(GPI)-A | 23 | 5.41E-04 | 0.081 |
| SAB | Agnostic | KEGG HISTIDINE METABOLISM | 24 | 5.54E-04 | 0.082 |
| SAB | Agnostic | abnormal physiological response to xenobiotic | 402 | 5.91E-04 | 0.085 |
| SAB | Agnostic | regulation of stress-activated MAPK cascade | 140 | 5.94E-04 | 0.085 |
| SAB | Agnostic | Metabolism of amino acids and derivatives | 171 | 6.39E-04 | 0.089 |
| SAB | Agnostic | epithelial cell differentiation involved in kidney c | 20 | 6.46E-04 | 0.089 |
| SAB | Agnostic | regulation of stress-activated protein kinase sign | 141 | 7.20E-04 | 0.098 |
| SAB | Hypothesis driven | MORNING | 109 | 2.29E-04 | 0.018 |
| SAB | Hypothesis driven | LOW | 8153 | 2.53E-04 | 0.018 |
| SAB | Hypothesis driven | Mitochondrion_(core) | 174 | 2.94E-04 | 0.018 |
| SAB | Hypothesis driven | ID-NS | 116 | 5.62E-04 | 0.025 |

We found significant enrichments between our BD associated genes and GWAS-derived gene sets for schizophrenia (p= 3.69E-13; all p-values shown are FDR-corrected), bipolar disorder (p= 2.59E-09) and major mood disorder (p=0.0040). These results are reassuring rather than illuminating, given the known genetic overlap between these disorders, the likely shared samples with the previous BIP GWAS, and the potential for shared controls between all PGC GWAS studies. Similar to the BD results, BD-1 associated genes were significantly enriched for GWAS-derived SCZ (p= 5.39E-12) and BD (p= 1.78E-09) gene sets. BD-II associated genes were not significantly enriched with previous BP or schizophrenia GWAS results. SAB-associated genes were significantly enriched with bipolar GWAS results (p= 0.027).

We identified three drug target gene sets enriched in our BPD associated genes; anabolic steroids (p=5.84E-4), androgens (p=0.025) and corticosteroids for systemic use (p=0.012). Corticosteroids when given in high doses can cause symptoms of mania, psychosis, impulsivity, irritability, anxiety, and depression^56,57^.

Four pathways in our ‘hypothesis-driven’ analysis were associated with BPD after FDR correction, including genes associated with self-defined ‘morning person’ chronotype^58^, genes that were highly intolerant to deleterious mutation in EXAC, genes with non-synonymous mutations linked to schizophrenia, and targets of the FMRP protein. FMRP pathways have previously been associated with schizophrenia, autism, and intellectual disability^33,59,60^. We identified five further pathways with nominally significant competitive MAGMA p-values, but which did not survive FDR-correction, relating to pre- and post-synaptic density, circadian clock genes, and loss of function mutations associated with intellectual disability.

For BD-I, we identified two associated pathways in the hypothesis-driven analysis after FDR correction; endoplasmic reticulum function (ER; p=0.036) and post synaptic density (PSD; p=0.046). 49/8,500 molecular pathways from public databases were significant after FDR-correction, with the most significant driven by methyltranferase activity (S-adenosylmethionine - dependent methyltransferase activity; p=3.0x10^−3^). Four pathways involved in methyltransferase activity are driven by TFB1M, a brain-expressed mitochondrial methyltransferase gene involved in neurosensory mitochondrial deafness^61,62^. Other significant pathways include mitochondrial function (mitochondrial genome maintenance; p=0.032) which was also validated in studies of the PSD proteins and associations with bipolar disorder^63^.

For BD-2 there were no significant hypothesis-driven pathways; however, 34 agnostic pathways were significantly enriched. S-adenosylmethionine-dependent methyltransferase activity pathway was the most significant (p=0.0029), in line with our BD-I analysis. Other significant pathways and potentially interesting pathways include metabolism of porphyrins, heme biosynthesis, abnormal neuronal migration, and negative regulation of systemic arterial blood pressure.

Three hypothesis-driven pathways were enriched with SAB; including mitochondrion^64^, non-synonymous mutations associated with intellectual disability, and genes that have low-level intolerance to EXAC mutations. Our large agnostic analysis revealed many neuron specific genes sets including axonal regeneration, Schwann cell differentiation, and neuron projection regeneration. Mitochondrion and mitochondrion localization were also significant further emphasizing the involvement of mitochondrial genes in bipolar disorder^65–67^. A total of 45 pathways were significantly enriched after FDR correction.

## Discussion

In this study, we present the largest analysis to date of transcriptomic imputation in Bipolar Disorder, and three bipolar disorder subtypes. Transcriptomic Imputation approaches leverage carefully curated eQTL reference panels to create prediction models of genetically-regulated gene expression^28,32,33,68^ (GREX). These models are then used to predict GREX in genotyped samples (for example, those obtained through GWAS), thus providing large, well-powered gene-expression datasets, while circumventing the difficulties and complications inherent in traditional transcriptome studies.

We applied gene expression predictor models derived from GTEX and CMC data to 21,488 bipolar disorder cases and 54,303 controls from the PGC-BD and iPSYCH collections, and obtained predicted genetically regulated gene expression levels (GREX) for 19,661 unique genes, across 13 brain regions. We identified 53 independent BPD gene-tissue associations; of these, 29 were novel, i.e., they did not occur within 1MB of a locus identified in the recent PGC-BD GWAS^5^. Additionally, we identified 46 independent subtype-specific gene-tissue associations.

Our study includes an additional 1,503 BPD cases and ~23,000 controls from the iPSYCH consortium, which were not included in the discovery stage of the recent PGC-BD GWAS, and so some proportion of these novel associations likely stem from both the increased power of our sample, as well as the increased power of prediXcan over GWAS^28,33^. It should be noted that our BD-II, SAB, and cross-subtype analyses are small, and power to detect true associations is therefore low. These analyses should be taken as preliminary, exploratory findings, and larger, more well-powered studies should be carried out.

BPD- and BD-I-associated genes identified in this study were significantly more likely to be differentially expressed in post-mortem tissue from individuals with bipolar disorder than might be expected by chance. Replication of highly associated genes was tissue-specific; for example, genes discovered in the DLPFC were differentially expressed in the DLPFC. When testing only nominally significant genes (i.e., all genes reaching p<0.05), replication was highly similar across all tissues, and degree of replication seemed to be driven by the power of the original eQTL reference panel (taking sample size as a proxy). This might indicate a large group of genes with broad, multi-region implications, while smaller groups of genes confer region-specific BPD risk.

It is likely that some of the cross-brain signal also arises from highly correlated gene expression patterns and shared eQTLs between brain regions^32,55^. We used microarray data from a small sample of individuals with BD-II to visualize expression of our two BD-II associated genes,

*NUP98* and *COLGALT1*, in cases compared to controls. For both genes, the observed direction of effect matches our prediXcan results. Although these results are encouraging, this analysis is based on a very small number of cases; as such, these results should be interpreted as early, preliminary indications, which should be followed with larger and more detailed investigations.

An interesting feature of transcriptomic analysis is the ability to probe associations across specific brain regions (Suppl. Table 1). In our BPD meta-analysis, we identified 20 pre-frontal cortex associations (nine in the DLPFC), 13 in the striatum (Caudate, Nucleus Accumbens, and Putamen Basal Ganglia), 11 in the cerebellum and cerebellar hemisphere, and 2 in the hippocampus. These results imply prominent roles for the frontal cortex, striatum and cerebellum in bipolar disorder, consistent with previous neuro-anatomical studies. For example, imaging studies have repeatedly demonstrated enlarged putamen^69–71^ and caudate^69,72–74^ regions, decreased cerebellar volumes^69,75–77^, and structural differences in the prefrontal cortex of individuals with BPD^69,78–81^.

We used genic associations for BD, BD-I, BD-II, and SAB to search for pathway enrichment with MAGMA^42^ using gene sets for drug targets, hypothesis driven, and agnostic gene sets. Our drug target genes revealed sets for anabolic steroids, corticosteroids, and androgens which have common precursors and similar effects on hormone receptors. Hormone imbalance has been hypothesized in patients with BD and schizophrenia. Altered hypothalamic-pituitary-adrenal (HPA) axis and increased systemic cortisol metabolism was found by measuring cortisol metabolizing enzymes in urine of patients vs controls suggesting the synthesis pathways for these hormones are altered^57^. Corticosteroids themselves are prescribed for a number of different medical conditions and can cause symptoms in patients that include psychosis, mania, depression, mixed features, delirium, and anxiety^82^. While these symptoms can arise after corticosteroid use, we cannot be certain the mechanisms are unique and the shared phenotypes in these overlapping gene sets suggest a similar genetic underpinning. Further investigation is warranted to understand the pathways involved in corticosteroid induced psychiatric symptoms and symptoms experienced by patients in bipolar disorder and schizophrenia. Additionally, our pathway analysis results provide support for a number of specific biological hypotheses.

### Oxidative Stress and Mitochondrial Dysfunction

Collectively, our results indicate a potential role for oxidative stress and mitochondrial dysfunction in bipolar disorder. This hypothesis has been explored in detail elsewhere^83–86^, and has been implicated in BPD ^83–85^ as well as a range of psychiatric disorders^87–90^, including anxiety and panic disorders^91^, schizophrenia^92–94^, and major depressive disorder^95^. Evidence for the involvement of oxidative stress and mitochondrial dysfunction in BPD includes known comorbidities between bipolar disorder and mitochondrial disease^96^, the known antioxidant properties of antipsychotic drugs^83^, and the demonstrated benefit of antioxidant therapies in individuals with schizophrenia and bipolar disorder^83^.

A substantial number of the genes identified in our meta-analyses also have a role in oxidative stress and mitochondrial dysfunction (including for example, *AIFM3, CHDH, EDEM2, EIF1AD, FADS1, TARS2).* In particular, our PHEWAS results implicate a gene, *EIF1AD*, which has a wel-described role in response to oxidative stress^97^. Reduced expression of *EIF1AD* (eukaryotic translation initiation factor 1A domain containing; also known as haponin) in the DLPFC was associated with panic attacks, mixed states, and BD-I; in line with this, a recent study found increased RNA damage due to oxidative stress in individuals with BD-I and mixed states, compared to controls, and a decrease in levels of RNA damage after remission from an episode^84^. A large number of associations in our pathway analyses (Table 5) also point to mitochondrial methyltransferase pathways, endoplasmic reticulum function, mitochondrial function, and mitochondrion location.

Common with BD-I and BD-II are the methyltransferase pathways with the most significant genes involved in mitochondrial methyltransferase. These genes are responsible for neurological phenotypes and associated with bipolar disorder^65,66^. A study of human induced pluripotent stem cells found early mitochondrial abnormalities in lithium responsive patients with bipolar disorder suggesting these mitochondrial abnormalities are present at the earliest stages of cell development^67^. SAB significant pathways reinforce the relationship between bipolar disorder with mitochondrial and neuronal function.

### Post-synaptic Density

Multiple studies and hypotheses have implicated the post-synaptic density (PSD) as having a role for Bipolar Disorder, Schizophrenia, and other psychiatric disorders^63,64^. The PSD is a key location for a host of dopamine and glutamate signaling interactions, and has a key role in axonal growth and guidance. Further, proteins located in the PSD are involved in NMDA receptor trafficking, and underlie energy pathways and mitochondrial function. Our BD-I results are significantly enriched for genes related to PSD-95, a scaffolding protein within the PSD (p=5.2e-04). This enrichment is not driven by a single highly associated gene, but rather a large number of sub-threshold associations. The most significant post synaptic density (PSD) gene PACS1 (p=5.57e-05) codes for MHC-1 removal of membrane proteins in the trans golgi network and is overexpressed in brain; other subthreshold PSD-95 and glutamatergic associations include *TUBA1B* (p=3.1e-04), *SHANK1* (p=5.4e-04), *BSN* (p=6.5e-04), and *AP2B1* (p=6.7e-04). Additionally, our results are enriched for targets of the FMRP (fragile-X mental retardation protein; p=0.0015), in line with previous studies of Bipolar Disorder and schizophrenia^59,98^, as well as the original CommonMind Consortium analysis^31^. FMRP is encoded by *FMR1*, which is required at synapses for normal glutamate receptor signaling^99^.

### Circadian Rhythms

Longstanding hypotheses implicate the disruption of circadian rhythms in bipolar disorder. In particular, sleep disruption is included among bipolar disorder diagnostic criteria and is cited as a particular concern for individuals with BPD. Addressing circadian rhythm disruption is a key factor in treatment of bipolar disorder^100,101^, and in identifying individuals at risk of relapse^102–106^. Even among healthy individuals, circadian entrainment and sleep patterns are deeply entwined with mood regulation^100,107–112^. These relationships have been discussed in detail elsewhere, including detailed discussions of plausible neurobiological mechanisms^100,113–126^. Consequently, studies of the genetics of bipolar disorder have included an emphasis on “clock” genes, i.e., genes involved in regulating circadian rhythmicity^100,125,127,128^, and the genetics of chronicity and sleep traits^124^.

Our BPD-association results include genes with a role in regulation of circadian rhythm; *CIART* (Circadian Associated Repressor Of Transcription), *CNNM4, ZSWIM3, RPRD2, TARS2, HSPD1, VPS45* and *PHLPP1*, as well as *ASCC3 ^129^,DUSP7, ITGA9, VPS4A, MAPRE2, RRP12* and *CSE1L*, associated with BD-I; and *NUP98*, associated with BD-II, as well as ~30 other sub-threshold associated circadian rhythm genes (p<1e-03), including genes identified in a recent GWAS of self-identified ‘morning-ness’. These ‘morning-ness’ genes constituted the most significantly enriched set in our hypothesis-driven pathway analysis (p=3.27e-05) within the full bipolar meta-analysis; additionally, we identified enrichments for circadian clock genes (p=0.012) and clock modulators (p=0.023), although these did not remain significant after FDR-correction. ‘Morning-ness’ genes were also enriched among SAB prediXcan associations (p=2.3e-04) and BD-I associations (p=0.0012), although the latter does not survive FDR-correction (p=0.069).

## Acknowledgements

Data were generated as part of the CommonMind Consortium supported by funding from Takeda Pharmaceuticals Company Limited, F. Hoffman-La Roche Ltd and NIH grants R01MH085542, R01MH093725, P50MH066392, P50MH080405, R01MH097276, RO1-MH-075916, P50M096891, P50MH084053S1, R37MH057881 and R37MH057881S1, HHSN271201300031C, AG02219, AG05138 and MH06692.

Brain tissue for the study was obtained from the following brain bank collections: the Mount Sinai NIH Brain and Tissue Repository, the University of Pennsylvania Alzheimer’s Disease Core Center, the University of Pittsburgh NeuroBioBank and Brain and Tissue Repositories and the NIMH Human Brain Collection Core. CMC Leadership: Pamela Sklar, Joseph Buxbaum (Icahn School of Medicine at Mount Sinai), Bernie Devlin, David Lewis (University of Pittsburgh), Raquel Gur, Chang-Gyu Hahn (University of Pennsylvania), Keisuke Hirai, Hiroyoshi Toyoshiba (Takeda Pharmaceuticals Company Limited), Enrico Domenici, Laurent Essioux (F. Hoffman-La Roche Ltd), Lara Mangravite, Mette Peters (Sage Bionetworks), Thomas Lehner, Barbara Lipska (NIMH).

The iPSYCH-GEMS team would like to acknowledge funding from the Lundbeck Foundation (grant no R102-A9118 and R155-2014-1724), the Stanley Medical Research Institute, an Advanced Grant from the European Research Council (project no: 294838), the Danish Strategic Research Council the Novo Nordisk Foundation for supporting the Danish National Biobank resource, and grants from Aarhus and Copenhagen Universities and University Hospitals, including support to the iSEQ Center, the GenomeDK HPC facility, and the CIRRAU Center.

The Genotype-Tissue Expression (GTEx) Project was supported by the <u>Common Fund</u> of the Office of the Director of the National Institutes of Health, and by NCI, NHGRI, NHLBI, NIDA, NIMH, and NINDS. The data used for the analyses described in this manuscript were obtained from the <u>GTEx Portal</u> on 09/05/16. BrainSpan: Atlas of the Developing Human Brain [Internet]. Funded by ARRA Awards 1RC2MH089921-01, 1RC2MH090047-01, and 1RC2MH089929-01.

---

**Supplementary Figure 1:**
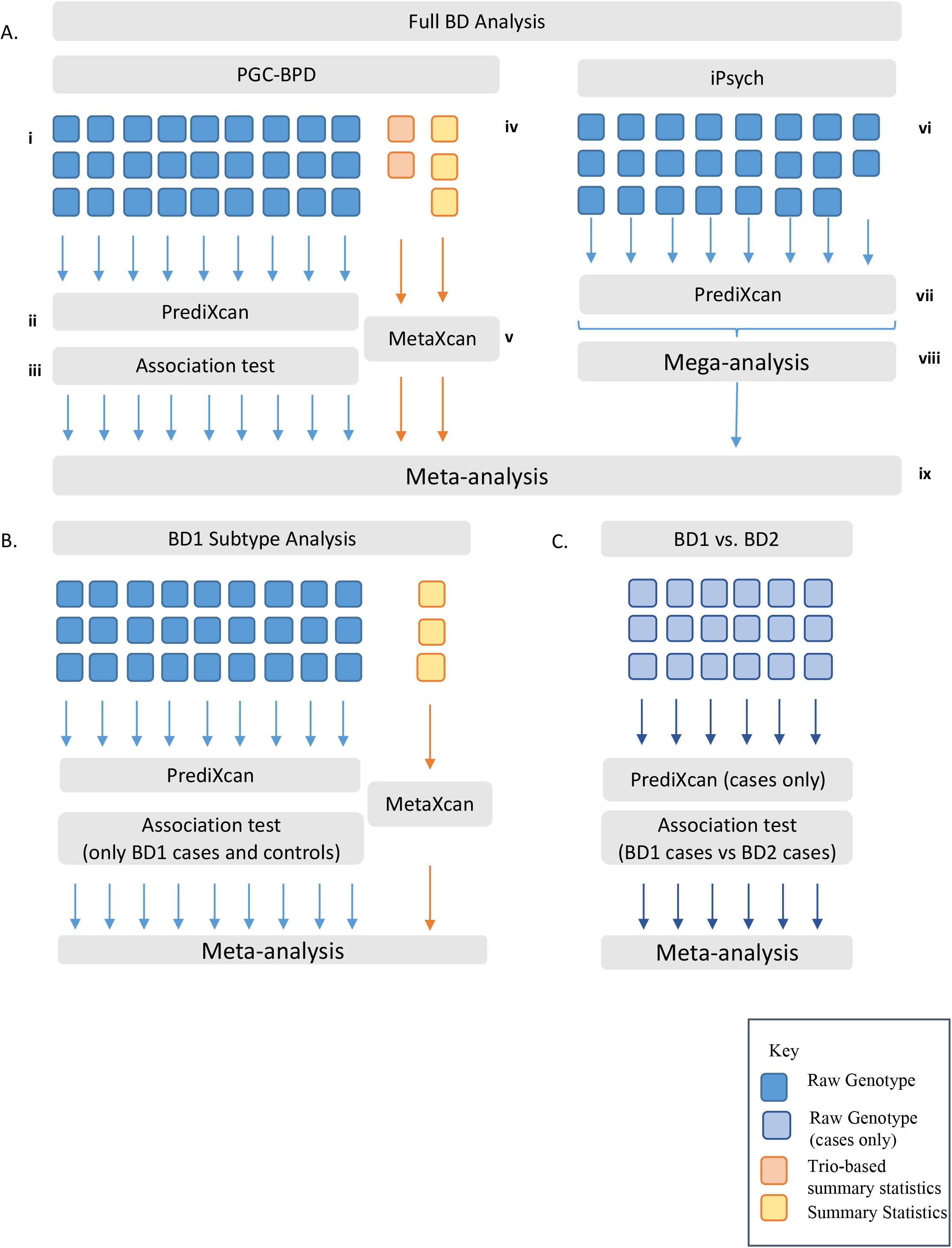
Analysis outline. A) Discovery Samples. 27 PGC-SCZ cohorts had available raw genotypes (i). Predicted DLPFC gene expression was calculated in each cohort using prediXcan (ii) and tested for association with case-control status (iii). 5 PGC cohorts (2 trio, 3 case-control) had only summary statistics available (iv). MetaXcan was used to calculate DLPFC associations for each cohort (v). iPsych samples were collected in 23 waves (vi). Predicted DLPFC gene expression was calculated in each wave separately using prediXcan (vii) and merged for association testing. A mega-analysis was run across all 23 waves, using wave membership as a covariate in the regression (viii). Results were meta-analysed across all 32 cohorts and the iPsych MEGA-analysis results(ix). This procedure was repeated for 12 GTEx prediction models. B) Subtype Analyses. Subtype information was available only for PGC-BD samples. Analysis was carried out in the same way as for the full BD analysis (A), including only BD1 cases. C) Cross-subtype analysis. Analysis was carried out for cases only, in the same way as A andB.

